# Evolution of diversity in metabolic strategies

**DOI:** 10.1101/2020.10.20.347419

**Authors:** R. A. Caetano, Y. Ispolatov, M. Doebeli

**Author notes:** **For correspondence:** (FMS).

## Abstract

Understanding the origin and maintenance of biodiversity is a fundamental problem. Many theoretical approaches have been investigating ecological interactions, such as competition, as potential drivers of diversification. Classical consumer-resource models predict that the number of coexisting species should not exceed the number of distinct resources, a phenomenon known as the competitive exclusion principle. It has recently been argued that including physiological tradeoffs in consumer-resource models can lead to violations of this principle and to ecological coexistence of very high numbers of species. Here we show that these results crucially depend on the functional form of the tradeoff. We investigate the evolutionary dynamics of resource use constrained by tradeoffs and show that if the tradeoffs are non-linear, the system either does not diversify, or diversifies into a number of coexisting species that does not exceed the number of resources. In particular, very high diversity can only be observed for linear tradeoffs.

## Introduction

Life on Earth is spectacularly diverse ***May (1988)***. For example, one study in the early 2000’s found that the number of species of fungi is, by a conservative estimate, ca. 1.5 M ***Hawksworth (2001)***, which was subsequently revised to be between 2.2 to 3.8 M species ***Hawksworth and Lücking (2017)***. Microbes are by far the most diverse form of life. They constitute approximately 70-90% of all species ***Larsen et al. (2017)***. Perhaps even more astonishing than the number of species is the fact that all of them came from a single common ancestor ***Darwin (1859)***; ***Steel and Penny (2010)***; ***Theobald (2010)***. To understand the fundamental mechanisms behind such diversification is one of the most relevant problems addressed by the scientific community ***Mayr and Mayr (1963)***; ***Coyne (1992)***; ***Rice and Hostert (1993)***; ***Higashi et al. (1999)***; ***Dieckmann and Doebeli (1999)***; ***Gavrilets and Waxman (2002)***; ***de Aguiar et al. (2009)***.

Recently, ecological interactions, such as competition, have received a lot of attention as potentially very strong drivers of diversification and speciation. A widely used class of models in which this phenomenon can be observed is based on classical Lotka-Volterra competition models, which are augmented by assuming that the carrying capacity is a (typically unimodal) function of a continuous phenotype, and that the strength of competition between two phenotypes is measured by a competition kernel, which is typically assumed to be a (symmetric) function of the distance between the competing phenotypes, with a maximum at distance 0 (so that the strength of competition decreases with increasing phenotypic distance).

These assumptions are biologically plausible, and such models have been widely used to provide insights into evolutionary diversification due to competition ***Dieckmann and Doebeli (1999)***; ***Doebeli and Ispolatov (2010***, 2017). However, these models are not derived mechanistically from underlying resource dynamics, and in fact it is known that the commonly used Gaussian functions for the carrying capacity and the competition kernel are not compatible with resource-consumer models ***Abrams (1986)***; ***Ackermann and Doebeli (2004)***. A more mechanistic approach is desirable.

Recently, a MacArthur consumer-resource model ***Macarthur and Levins (1967)*** was studied in an ecological context with a view towards explaining the existence of very high levels of diversity ***Posfai et al. (2017)***; ***Erez et al. (2020)***. The authors consider different species competing for R interchangeable resources, each supplied at a constant rate ***Posfai et al. (2017)*** or periodically repleted after being used ***Erez et al. (2020)***. A consumer species is characterized by an uptake strategy, *α* = (*α*_1_, …, *α*_*R*_), where the *j* th component *α*_*j*_ ≥ 0 represents the amount of cellular metabolism allocated to the uptake of the *j* th resource. The total amount of cellular metabolism available for resource uptake is limited, and hence it is natural to assume a tradeoff between the uptake rates of different resources. In general mathematical terms, a tradeoff is typically given by a function *T* (*α*) = *T* (*α*_1_, …, *α*_*R*_) that is increasing in each of the arguments *α*_*j*_, and such that the only permissible allocation strategies *α* are those satisfying *T* (*α*) ≤ *E*, where *E* is a constant. The analysis is then typically restricted to the subspace of strategies defined by *T* (*α*) = *E* (because *T* is increasing in each *α*_*j*_). It was shown in ***Posfai et al. (2017)***; ***Erez et al. (2020)*** that, under the assumption of a linear tradeoff, 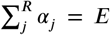, very high levels of diversity, i.e., many different species with different *α*-strategies, can coexist. This is a very interesting finding because it violates the competitive exclusion principle ***Hardin (1960)***, according to which at most *R* different species should be able to stably coexist on *R* different resources. Such high levels of diversity emerging from simple consumer-resource models could help solve the paradox of the plankton ***Hutchinson (1961)*** from an ecological perspective.

Here, we incorporate evolutionary dynamics into the ecological models of ***Posfai et al. (2017)***; ***Erez et al. (2020)*** and investigate the conditions under which diversity evolves from single ancestors. To make the model more general and relevant, we consider non-linear tradeoffs in resource use. The non-linearity of tradeoffs is a direct and inevitable consequence of general non-linearity of chemical kinetics and equilibria. The rate and mass-action equilibrium of even a simple bimolecular reaction are non-linear functions of the concentrations of reactants. So probably the easiest way to see that tradeoffs between uptakes rates of different resources have to be non-linear as well is to assess physiological costs of enzymatic and transport machinery involved in metabolic functions. Such machinery always include hetero- and homo-oligomeric complexes, for which the scaling between the concentration of metabolically-active effectors complexes and their more elementary constituent parts is inevitably non-linear: Doubling the concentration of am oligomer requires more than doubling the concentrations of its monomers. So the metabolic cost of the former in the units of the metabolic costs of the later is non-linear as well. Besides this example, there are plenty other ways to convince oneself that physiological costs of maintaining a specific metabolic pathway is a non-linear function of the productivity of such a pathway. In this paper, we incorporate non-linear effects into the tradeoff function by considering the energy budget as a sum

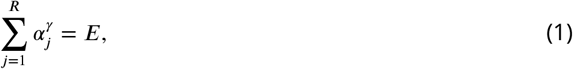

where *γ* and *E* are positive constants. We show that in the resulting evolutionary model, coexistence of more than *R* species only emerges for the (structurally unstable) linear case *γ* = 1. Using adaptive dynamics and numerical simulations, we show that regardless of the value of *γ*, an initially monomorphic population always evolves to an attractive fixed point (also called “singular point”), after which two generic scenarios are possible: *i)* if *γ* < 1, the population branches and diversifies, with the maximal number of coexisting species equal to the number of resources *R*, a state in which each species is a complete specialist on exactly one of the resources; *ii*) if *γ* > 1, an initially monomorphic population also evolves to a singular point, but subsequently does not diversify and instead remains a monomorphic generalist. To make the argument for the relevance of non-linear tradeoffs even more clear, we show that an omnipresent non-linearity in the dependence of nutrient uptake rates on *α* can be transformed into the non-linearity of tradeoff (1) and vice versa. So either non-linearity is sufficient to brings the diversity down to the competitive exclusion limit. We also show that two scenarios (of a generalist and *R* complete specialists) emerge as results of pure ecological dynamics in the system initially populated with multiple species with various uptake strategies *α* that satisfy (1).

Overall, our results show that very high levels of diversity do not evolve in the consumer-resource model considered here in a realistic scenario where tradeoffs in resource preference or the resource uptake rates are non-linear.

## Model and Results

We consider a population competing for *R* substitutable resources in well-mixed environments. A phenotypic species *α* is characterized by its metabolic allocation strategy *α* = (*α*_1_, …, *α*_*R*_), where *α*_*j*_ is the per capita rate at which individuals of species *α* take up the *j* th nutrient. Various coexisting species are distinguished by there specific *α*s. From a physiological perspective, *α*_*j*_ is proportional to the amount of metabolic effort allocated by the individuals of species *α* to capture nutrient *j*. Intrinsic limitations on metabolic activities impose a restriction on the total amount of nutrient up-take. For simplicity, we assume that this intrinsic limitation leads to a tradeoff in the components *α*_*j*_ of the form (1). (Note that we also assume *α*_*j*_ ≥ 0 for all *j*.) Throughout, we will set the scaling parameter *E* = 1. (See Appendix for a more general treatment, in which the exponent *γ* can differ for different directions *α*_*j*_ in phenotype space.)

Following ***Posfai et al. (2017)***, we denote by *c*_*j*_ (*t*) the concentration of resource *j* at time *t*, and we assume that the amount of resource *j* available for uptake per individual (e.g., the amount of resource bound to the outer membrane of a microbial cell) is given by a monotonously increasing function *r*_*j*_ (*c*_*j*_). Specifically, we assume this function to be of Monod type, *r*_*j*_ (*c*_*j*_) = *c*_*j*_ /(*K*_*j*_ + *c*_*j*_). Thus, the rate of uptake of resource *j* by an individual consumer with uptake strategy *α* is *α*_*j*_ *r*_*j*_ (*c*_*j*_).

### Chemostat conditions

We assume that resources are supplied to the system at a constant rate defined by the supply vector *S*= (*S*_1_, …, *S*_*R*_), so that resource *j* is supplied at a constant total rate *S*_*j*_ and decays at a rate *µ*_*j*_ ***Posfai et al. (2017)***. This generates the following system of equations for the ecological dynamics of the concentrations *c*_*j*_, *j* = 1, …, *R*:

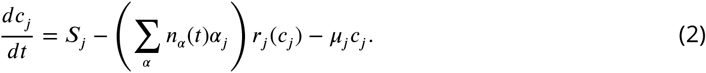

Here *n*_*α*_ (*t*) is the population density of species *α* at time *t*, so that ∑_*α*_ *n*_*α*_ (*t*) *α*_*j*_ is the total amount of metabolic activity invested into uptake of resource *j* (the sum runs over all species *α* present in the community). We further assume that the cellular per capita birth rate of species *α* is equal to the amount of nutrient absorbed by each individual. The dynamics of the population density *n*_*α*_ then becomes

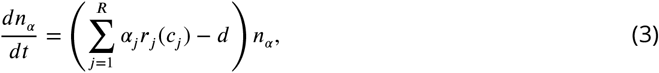

where *d* is the per capita death rate, which is assumed to be the same for all consumers.

The evolutionary dynamics of the the traits *α*_*j*_ can be solved analytically only for a simplified system in which the resource decay (dilution) rates *µ*_*j*_ are set to 0. This assumption, also made in ***Posfai et al. (2017)***, corresponds to rapid consumption of almost all resource. In the Appendix, we derive the adaptive dynamics for the allocation strategies, i.e., for the traits *α*_*j*_ (***Metz et al. (1992)***; ***Dieckmann and Law (1996)***; ***Dieckmann and Doebeli (1999)***; ***Doebeli and Dieckmann (2003***, 2000); ***Hui et al. (2018)***; ***Doebeli (2011)***; ***Geritz et al. (1997)***). We show that with vanishing decay rates, there is a unique singular point

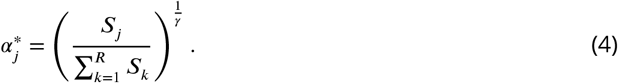

Calculations of the Jacobian of the adaptive dynamics (an indicator of convergence stability of a fixed point) and of the Hessian of the invasion fitness function (which distinguishes whether the fixed point is an evolutionary endpoint or a branching point) yield the following conclusions: Regardless of the value of *γ*, the singular point *α*^∗^ is always convergent stable, so that the system approaches *α*^∗^ from any initial condition. If *γ* > 1, the singular point *α* is also evolutionarily stable and hence represents the evolutionary endpoint. In particular, no diversification takes place. On the other hand, if *γ* < 1, the singular point is evolutionarily unstable, and hence is an evolutionary branching point. In particular, if *γ* < 1, the system will diversify into a number of coexisting consumer species. If *γ* = 1 (linear tradeoff), the fitness Hessian is 0, representing evolutionary neutrality.

To check our analytical approximations and to investigate the details of diversification after convergence to the evolutionary branching point, we performed numerical simulations of evolving populations consisting of multiple phenotypic strains. The simulations were performed without the simplifying assumption of zero resource degradation (dilution) rates, further details of the numerical simulations are presented in the Appendix.

In the figures below we show evolving populations as circles with radii proportional to the square root of population size *n*_*α*_ in 3-dimensional strategy space (*α*_1_, *α*_2_, *α*_3_), viewed orthogonally to the simplex plane 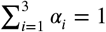. With the constraint 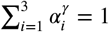, the coordinates of each population are 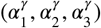. In the following numerical examples we considered a symmetric supply of resources ***S***_*i*_ = 1 and a slow resource degradation, *µ*_*i*_ ***K***_*i*_ = 0.1.

We first consider scenarios with linear tradeoffs, *γ* = 1. Figure 1 shows the evolution of a population (shown in blue circles) whose individuals die at constant rate *d* = 1 (corresponding videos of the simulations can be accessed through the links provided in the figure legends). The black circle represents the singular point that is calculated in the limit of low degradation of nutrients, given by Eq. (4). Figure 1 (a) shows the initial monomorphic population far from the singular point. An intermediate time of the evolutionary process is shown in Figure 1 (b), in which the population remains monomorphic and is approaching the singular point *α*^∗^. For *γ* = 1, the singular point is neutral evolutionarily (all eigenvalues of the Hessian of the invasion fitness function are 0 due to the linearity of the tradeoff), and once the population converges to the singular point, it starts to diversify “diffusively”, as anticipated in ***Posfai et al. (2017)***: neutrality of selection results in communities consisting of a large number of species. Thus, the high diversity observed in this case is an evolutionary consequence of the selective neutrality caused by a linear enzymatic trade-off.

**Figure 1.**
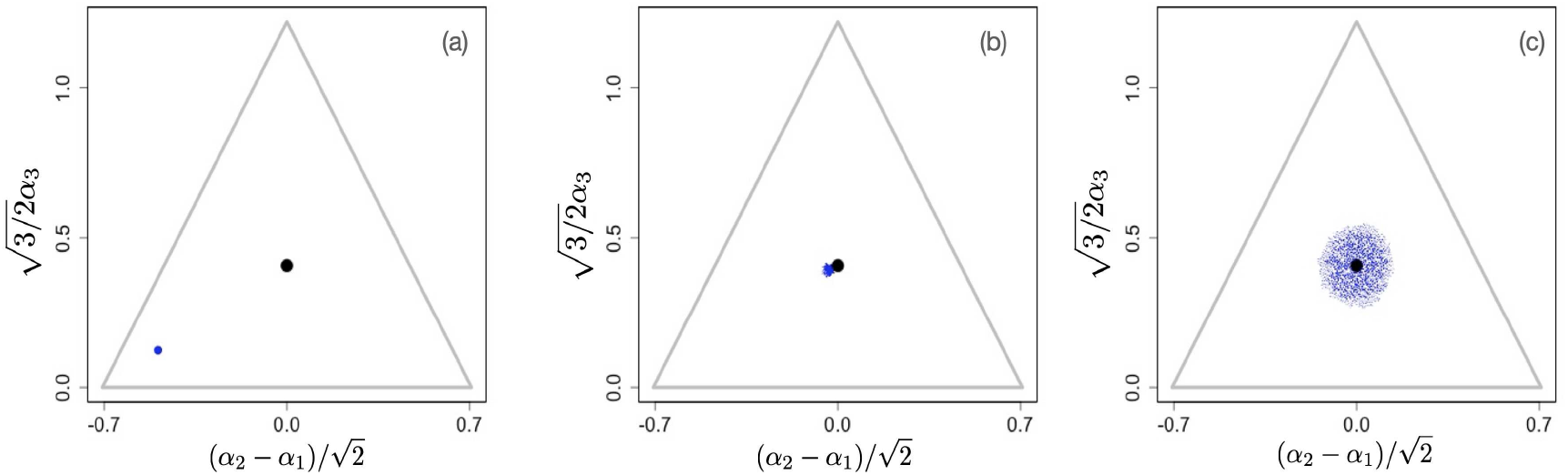
Snapshots illustrating the beginning, intermediate, and advanced stages of evolution under a linear constraint, *γ* = 1. A video of the entire evolutionary process can be found **[here](https://figshare.com/s/d59cf5c2f344424f830a)**, frames are recorded every 200 time units until *t* = 30000 and then, to better illustrate slow neutral evolution, the frame recording times *t*_*i*_ were defined as a geometric progression *t*_*i*+1_ = 1.006*t*_*i*_. Other parameter values were *S*_*j*_ = 1, *µ*_*j*_*K*_*j*_ = 0.1 for *j* = 1, 2, 3, and *d* = 1,

The situation changes for non-linear tradeoffs, *γ* ≠ 1, which generates two very different evolutionary regimes depending on whether *γ* > 1 or *γ* < 1 (even when the deviation of *γ* from one is small). Figure 2 (a-c) shows an example of the evolutionary dynamics for *γ* = 1.1.

**Figure 2.**
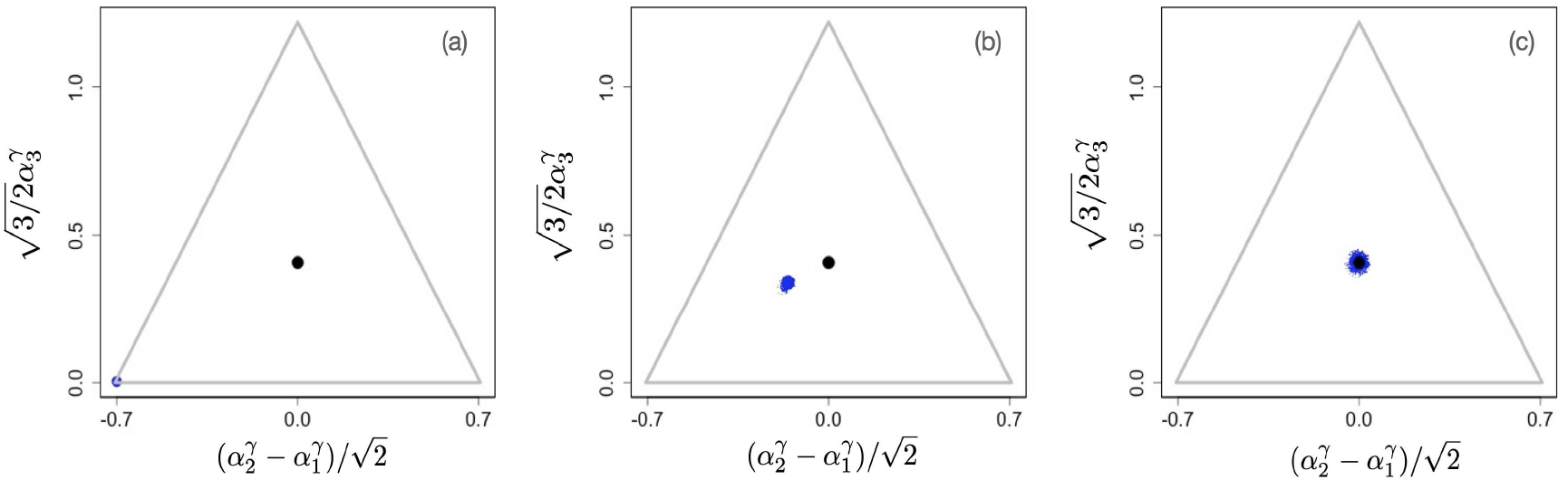
Example of evolutionary dynamics for *γ* = 1.1, showing convergence to the singular point given by Eq. (4) (and indicated by the black dot), but no subsequent diversification. The corresponding video can be found **[here](https://figshare.com/s/d59cf5c2f344424f830a)**, each frame in the video is separated by 1000 time steps. Other parameter values were *S*_*j*_ = 1, *µ*_*j*_*K*_*j*_= 0.1 for *j* = 1, 2, 3, and *d* = 0.25

The dynamics starts with an initial monomorphic population far from the singular point, as shown in Figure 2 (a). As in the linear case, and as predicted by the analytical theory, the monomorphic population converges toward the singular point Figure 2 (b). However, because *γ* > 1 the singular point is evolutionarily stable, and no diversification occurs (apart from mutation-selection balance around the singular point). Instead, when the population reaches the singular point, evolution comes to a halt, and all individuals are generalists, i.e, use all resources to some extent (as determined by the location of the singular point), as depicted in Figure 2 (c).

On the other hand, Figure 3 (a-c) shows the evolutionary process for a community with *γ* = 0.9. The initial configuration is shown in Figure 3 (a). As in the previous examples, the initial phase of evolution ends with the population converging to the singular point *α*^∗^. However, in this case, the singular point is an evolutionary branching point giving rise to the emergence of distinct and diverging phenotypic clusters (Figure 3 (b)). The final state of the evolutionary process is shown in Figure 3 (c): there are three coexisting phenotypic clusters, each being a specialist in exactly one of the resources. Our numerical simulations indicate that the results shown in Figs. 1-3 are general and robust: non-neutral diversification occurs only for *γ* < 1 and typically leads to coexistence of *R* specialists. In fact, the results easily generalize to situations in which the exponent *γ* in the tradeoff function may be different for different directions in phenotype space, i.e., for different *α*_*j*_. As we show in the Appendix, evolutionary branching along a direction *α*_*j*_ in phenotype space can occur if the corresponding exponent *γ*_*j*_ < 1. Figures 1 and 2 in the Appendix illustrate scenarios in which only a subset of the phenotypic directions *α*_*j*_ are branching directions along which evolutionary diversification occurs. In such a case, the number of distinct species resulting from the evolutionary process is less than *R*.

**Figure 3.**
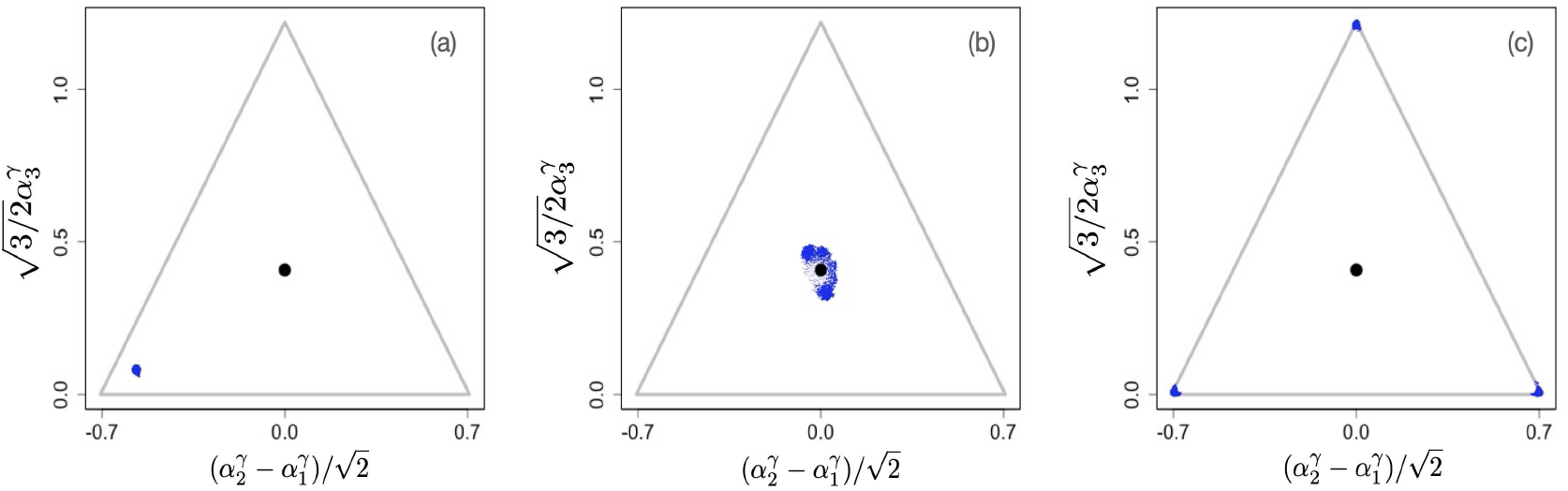
Example of evolutionary dynamics for *γ* = 0.9, showing initial convergence to the singular point (indicated by the black dot) and subsequent diversification into three specialists, each consuming exclusively one of the three resources. The corresponding video can be found **[here](https://figshare.com/s/d59cf5c2f344424f830a)**, each frame in the video is separated by 1000 time steps. Other parameter values were *S*_*j*_= 1, *µ*_*j*_*K*_*j*_ = 0.1 for *j* = 1, 2, 3, and *d* = 0.25,

Finally, we note that our results for the effects of non-linear tradeoffs on evolutionary dynamics have corresponding results in purely ecological scenarios, such as those studied in ***Posfai et al. (2017)***. We simulated ecological time scales by seeding the system with a set of e.g. randomly chosen phenotypes throughout phenotype space and running the population dynamics with the mutational process turned off. Again, as shown in Appendix 1 Fig. 3, non-linear tradeoffs have a profound effect on the number of surviving species in such ecological simulations, with many species coexisting when *γ* = 1, as reported in ***Posfai et al. (2017)***, but with typically only *R* species surviving when *γ* < 1 and only very few species surviving in the close vicinity of the singular point when *γ* > 1.

### Serial dilution conditions

Serial dilution conditions are defined as a sequence of explicitly non-stationary inoculation and growth events ***Erez et al. (2020)***, which mimics seasonality or batch culture experiments (e.g. ***Lenski and Tra (1994)***). Each growth phase starts with the introduction of a diluted collection of species from a previous batch

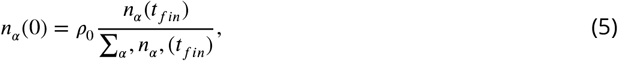

into a fresh batch of resources with a given composition *c*_*j*_ (0). In each batch, the species densities *n*_*α*_ (0) increase with time as

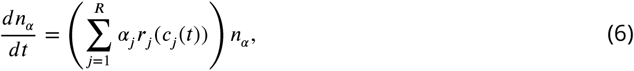

while resources are depleted:

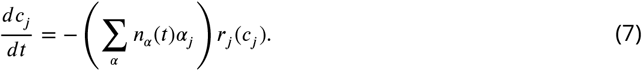

Unlike in the chemostat model, the death of individuals and the decay of resources are ignored (*d* = 0 and *μ* = 0). Each event ends at time *t*_*fin*_ when all resources are almost completely depleted,

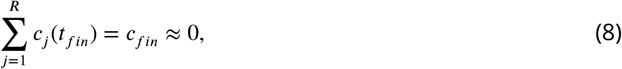

and the process is repeated.

Due to the explicit non-stationarity of such serial dilution processes, one of the main assumption of our adaptive dynamics analysis, the stationarity of resident populations, is not satisfied. Nevertheless, our numerical simulations show that the conclusions drawn for the chemostat case also hold for the serial dilution conditions, to the point that the simulation snapshots are visually indistinguishable from those shown in Figs. 2, 3. However, in the videos, which can be found **[here](https://figshare.com/s/3eb51bad58ae95b8b0f3)**, it is possible to see the oscillating population density, caused by the serial dilution protocol.

Specifically, we simulated the serial dilution for three limits considered in ***Erez et al. (2020)***, *c*_*j*_ (0) = 10*K, c*_*j*_ (0) = *K*, and *c*_*j*_ (0) = 0.1*K* for *ρ*_0_ = 10^−3^ and *c*_*fin*_ = 10^−8^. All other parameters were the same as used in Figs. 1, 2, 3 and corresponding videos.

In all three cases *c*_*j*_ (0) ≫ *K, c*_*j*_ (0) ∼ *K*, and *c*_*j*_ (0) ≪ *K* we observed that for *γ* > 1, the monomorphic population converges toward the singular point *α*^∗^ (Figure 2 (b) and video files **[here](https://figshare.com/s/3eb51bad58ae95b8b0f3)**). The singular point is evolutionarily stable, hence, as shown in Figure 2 (c), no subsequent diversification occurs (apart from narrow mutation-selection spreading around the singular point).

On the contrary, Figure 3 (a-c) and videos accessible **[here](https://figshare.com/s/3eb51bad58ae95b8b0f3)** show the evolutionary process for a community with *γ* < 1. The initial configuration is shown in Figure 3 (a). As in the previous examples, in the initial phase the monomorphic population evolves close to the singular point *α*^∗^. However, in this case, the singular point is an evolutionary branching point giving rise to the emergence of distinct and diverging phenotypic clusters (Figure 3 (b)). The final state of the evolutionary process is shown in Figure 3 (c): there are three coexisting phenotypic clusters, each being a specialist on one of the resources.

In addition, purely ecological (i.e., mutationless) simulations performed similarly to what is described above and in ***Erez et al. (2020)*** resulted in similar outcomes as in the chemostat model. In a system initially filled with many (200) species, only a few species survive after a fairly short transitory time. When *γ* > 1, 1 or a few species remain very close to the fixed point *α*^∗^, while for *γ* < 1, typically *R* specialist species remain in the system. The videos of pure ecological simulations can be seen **[here](https://figshare.com/s/3eb51bad58ae95b8b0f3)**)

## Discussion

To understand the origin and maintenance of diversity is a fundamental question in science. In particular, the mechanisms of diversification due to ecological interactions still generate lively debates.

Recently, tradeoffs in the rates of uptake of different resources were suggested as a mechanism to generate large amounts of diversity ***Posfai et al. (2017)***; ***Erez et al. (2020)***, possibly solving the “paradox of the plankton” ***Hutchinson (1961)***, and violating the competitive exclusion principle ***Hardin (1960)***, which states that the number of coexisting species should not exceed the number of resources. It has been shown that enzymatic allocation strategies that are plastic instead of fixed, so that individuals can change their allocation (while maintaining a linear tradeoff under a fixed allocation budget) in response to resource availability during their life time, tend to reduce the amount of diversity maintained in the ecological communities ***Pacciani-Mori et al. (2020b)***. Perhaps this is not surprising, since more plastic strategies tend to be able to be more generalist as well. As in ***Posfai et al. (2017)***; ***Erez et al. (2020)***, here we consider the case of non-plastic strategies, in which each individual is defined by its allocation vector *α*, but assuming a more general, nonlinear form of tradeoffs. Moreover, we investigate evolutionary rather than just ecological dynamics to determine the conditions under which evolutionary diversification can occur. There are no true jacks-of-all trades in biology and tradeoffs are a ubiquitous assumption in evolutionary thinking and modeling. However, the cellular and physiological mechanisms that underly such tradeoffs are typically very complicated and the result of biochemical interactions between many different metabolic pathways. Attempts have been made to understand tradeoffs more mechanistically, particularly in microbes ***Litchman et al. (2015)***, but higher-level modeling efforts most often still require a mostly phenomenological approach to incorporating tradeoffs. In this paper we assumed that each of *R* resources is available to each microbial organism at a certain rate that depends on the resource concentration in the system. The microbe in turn is described phenotypically by the metabolic allocation strategy that defines its uptake of the available resources.

Without tradeoffs, and everything else being equal, the best strategy would be to allocate an infinite amount (or at least the maximal amount possible) of metabolic activity to every resource, a scenario that is generally unrealistic biologically. Rather, tradeoffs inherent to cell metabolism prevent such strategies. Formally, tradeoffs are given by one or more equations (or more generally inequalities) that the phenotypes of individuals have to satisfy. In our simplistic models, tradeoffs are determined by the parameter *γ*, which essentially describes the curvature of the tradeoff function, with the linear tradeoff *γ* = 1 being the threshold between concave (*γ* < 1) and convex (*γ* > 1) tradeoffs. Formally, linear tradeoffs are the simplest case, but there is no a priori general reason why tradeoffs should be linear. Even for a single resource, the metabolic energy required for a given amount of enzymatic allocation to resource uptake is a non-linear function of that allocation due to non-linearity of enzymatic kinetics and binding equilibria in the concentrations of enzyme building blocks and other participating molecules.

Our results show that generically, diversity only evolves with concave tradeoffs, and the number of coexisting species never exceeds the number of resources. Only in the structurally unstable linear case (*γ* = 1) is it possible for very high levels of diversity to evolve due to the cessation of selection at the evolutionary equilibrium. Any value of *γ* ≠ 1 precludes high amounts of diversity. Extensive numerical explorations revealed that these results are robust and qualitatively independent of particular parameter choices, such as the number of resources, or the dynamics of resource input.

Furthermore, in the Appendix we show that the originally non-linear tradeoffs can be made linear by re-defining uptake rates *α*_*i*_ (16), thus “transferring” the non-linearity to the birth rate functions (11). But the metabolic and nutrient uptake rates are themselves non-linear in the enzyme concentration. In the Appendix we sketch a derivation of kinetics of an enzymatic reaction in the general case (assuming the steadiness of the concentration of the enzyme-substrate complex, yet without the assumption that the enzyme concentration is negligible compared to that of substrate). It follows that metabolic rates are in general sub-linear in the concentration of enzymes, which is intuitively clear if the rate saturation in the limit of infinite enzyme concentration is considered. However, sublinear rates are not the only possible deviation from linearity: the formation of enzyme oligomers ***Marianayagam et al. (2004)*** and spatially organized complexes ***Schmitt and An (2017)*** are controlled by intrinsically nonlinear mass-action equilibria, thus making the enzymatic rates generally sigmoid functions ***Ricard and Noat (1986)*** of the amount and thus physiological costs of production of individual enzymes. Whether sub- or super-linear, any deviation of the growth rates from the linear form (11, 6) results in a revalidation of the competitive exclusion limit, similarly to non-linearity in tradeoffs. This serves as another indication that the linear tradeoffs coupled to linear metabolic rates is a biologically unrealistic and exceptional case, while generic non-linearities do not generate high levels of diversity, and instead the outcomes are in line with classical results about the evolution of resource generalists vs. resource specialists ***Ma and Levin (2006)***.

It is well-known that the shape of tradeoff curves is, in general, an important component in adaptive dynamics models ***Kisdi (2015***, 2006). In particular, studies of evolution of cooperation (e.g. ***Damore and Gore (2012)***; ***Archetti and Scheuring (2012)***) have stressed that the outcome of evolution is conditional on the curvature of the public good and cost functions and provided numerous biochemical reasons for non-linearity of metabolic rates in enzyme concentrations. Here we have shown the importance of the tradeoff curvature for the evolution and maintenance of diversity in a general consumer-resource model. Of course, many potentially important ingredients that could yet lead to high or low diversity in these models were not considered in the present work. For example, dynamic and optimal metabolic strategies ***Pacciani-Mori et al. (2020a)*** and cross-feeding have recently been suggested as factors that could potentially enable such diversity ***Goyal and Maslov (2018)***, while “soft constraints” that allow random deviations of metabolic strategies from the exact tradeoff constraint, were reported in ***Cui et al. (2020)*** to reduce the diversity even below the competitive exclusion limit. It will be interesting to consider these model extensions with non-linear tradeoffs.

## Apppendix 1

### Ecological and Evolutionary Dynamics

We assume as in ***Posfai et al. (2017)*** that metabolic reactions occur on a much faster time scale than cellular division, so that resource concentrations are always at their ecological equilibrium values 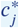 determined as solutions of equations

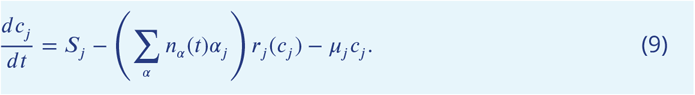

with *dc*_*j*_ /*dt* = 0. (Note that these equilibrium resource concentrations are determined by the current populations sizes *n*_*α*_ (*t*).) In practice, a faster time scale can be achieved by multiplying the right-hand side of (9) by a large dimensionless constant. The cellular per capita birth rate *g*_*α*_ of species *α* is proportional to the amount of nutrient absorbed by each individual,

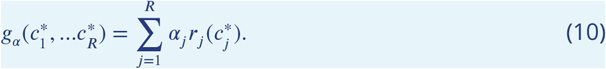

The dynamics of the population size *n*_*α*_ then becomes

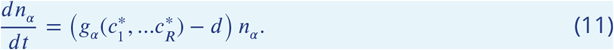

To derive the evolutionary dynamics for the allocation strategies, i.e., for the trait *α*, we follow the adaptive dynamics approach, a powerful tool to study gradual evolutionary diversification due to frequency-dependent ecological interactions ***Metz et al. (1992)***; ***Dieckmann and Law (1996)***; ***Doebeli (2011)***;***Dieckmann and Doebeli (1999)***; ***Doebeli and Dieckmann (2003***, 2000); ***Geritz et al. (1997)***; ***Hui et al. (2018)***. In particular, adaptive dynamics can generate the paradigmatic phenomenon of evolutionary branching ***Metz et al. (1992)***; ***Geritz et al. (1997)***; ***Doebeli and Dieckmann (2000)***; ***Hui et al. (2018)***; ***Doebeli (2011)***; ***Dieckmann and Doebeli (1999)***, during which a population that evolves in a continuous phenotype space first converges to a fitness minimum (evolutionary branching point) and then splits into two (or more) diverging phenotypic branches. We start with considering a monomorphic resident population at its ecological equilibrium 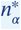, which is defined as the population size for which the equilibrium resource levels 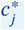 are such that 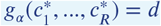 (note again that the 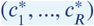 implicitly depend on *α*). The invasion fitness of a rare mutant *α*′ is then the per capita growth rate of the mutant *α*′ at the resource levels defined by the resident:

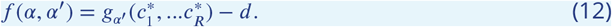

To derive the adaptive dynamics, we consider the selection gradient *q*(*α*) = (*q*_1_(*α*), …, *g*_*R*_(*α*)), with components

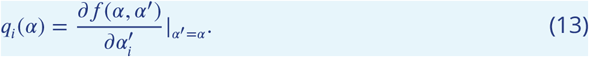

*g*(*α*) defines an *R*-dimensional dynamical system in unrestricted *α*-space,

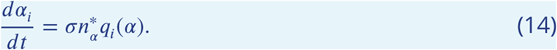

The speed of evolution of *α* is proportional to the current ecological equilibrium population size 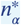 because the number of mutations occurring at any given point in time is proportional to 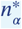. The parameter *σ* describes both the per capita rate and effective size of mutations. Without loss of generality, we set *σ* = 1.

To take the enzymatic tradeoff into account, the unconstrained adaptive dynamics (14) needs to be restricted to the surface in *α*-space that is defined by the tradeoff 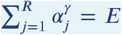, where *E* is a positive number ***Ito and Sasaki (2016)***. An illustrative example is the one in which the nutrients come from three different resources. The tradeoff 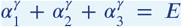 defines a surface in *α*-space where all strategies is embed in. The curvature of the surfaces are determined by *γ*. Figure 1 a) shows an example of the surface defined by the tradeoff for the case that *γ* > 1 while figures 1 b) and c) show the curvature for the case where *γ* = 1 and *γ* < 1, respectively. The blue star and the orange diamond illustrate possible position of the strategies in the *α* space. The individuals with strategy indicated by the blue star uptake nutrients only from resource *S*_1_ while the individuals with strategy indicated by the orange diamond uptake nutrients from the three resources.

**Appendix 1 Figure 1.**
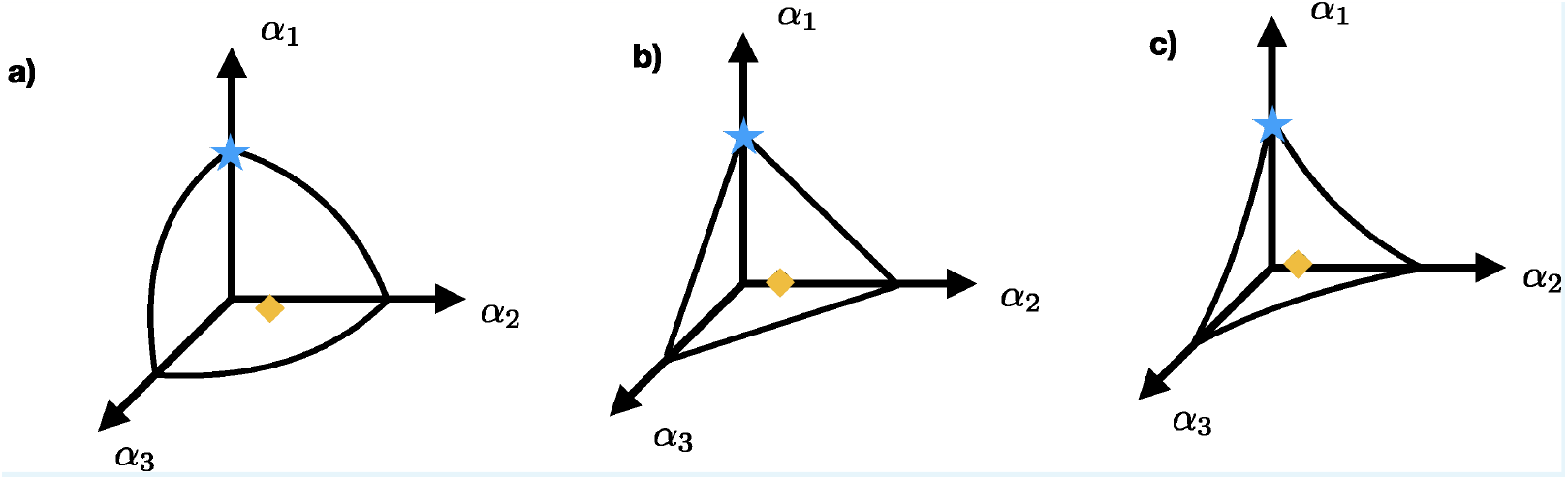
Three possible surfaces defined by tradeoff: a) shows the concave surface for the case where *γ* > 1 while b) and c) show the surface for the case where *γ* = 1 and *γ* < 1, respectively. The blue star and the orange diamond represent possible strategies. Individuals with strategy represented by the blue star obtain their nutrients only from resource *S*_1_ while the individuals that adopt strategy indicated by the orange diamond uptake nutrients from the three resources.

Equilibrium points of the adaptive dynamics, so-called singular points, are resting points *α*^∗^ of the resulting dynamical system in phenotype space. Given a singular point *α*^∗^, two stability concepts are important. First, there is stability in the usual sense of converging to *α*^∗^ from nearby initial conditions, which is measured by the Jacobian matrix of the functions defining the adaptive dynamics, evaluated at *α*^∗^. Second, evolutionary stability is measured by the Hessian of the invasion fitness function *f* (*α*^∗^, *α*′) with respect to the mutant trait and evaluated at the singular point *α*^∗^, and taken along the constraint surface ***Ito and Sasaki (2016)***. The negative definite Hessian (all eigenvalues being negative) means that the singular point is the maximum of invasion fitness and no branching occurs. Alternatively, a singular point is called an evolutionary branching point if it is both convergent stable with regard to the Jacobian and evolutionarily unstable with regard to the Hessian. Thus, a singular point is a branching point if all eigenvalues of the Jacobian have negative real parts, and if the Hessian matrix is not negative definite.

#### Singular points and their convergence and evolutionary stability

The case where the decay rates, *μ*_*j*_, are zero for any *j*, admits an analytical solution. We consider allocation strategies *α* = (*α*_1_, …, *α*_*R*_) as in the main text, but here we assume a more general tradeoff function:

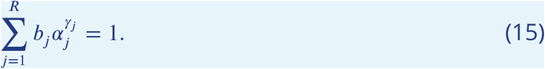

It turns out to be convenient to reparametrize the strategy space as follows:

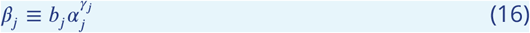

for *j* = 1, …, *R*. This simplifies the tradeoff expression to

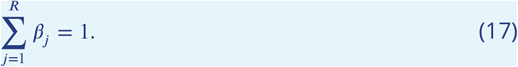

Because the *β*_*j*_ increase monotonically with *α*_*j*_, the adaptive dynamic properties in terms of convergence and evolutionary stability of singular points are the same for *α* = (*α*_1_, …, *α*_*R*_)- and *β* = (*β*_1_, …, *β*_*R*_)-phenotypes. However, the tradeoff in *β*, Eq. (17), is linear, which simplifies the analysis.

In terms of *β*, the per capita rate of use of resource *j* of an individual with phenotype *β* is

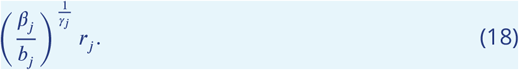

We assume that nutrients are supplied to the system at a constant rate given by the vector *S* = (*S*_1_, *S*_2_, …, *S*_*R*_), where *S*_*j*_ is the supply rate of the *j*th resource. We consider the low degradation rate regime, i.e., *μ*_*j*_ → 0 for all *j* in (9). Setting right hand sides of Eqs. (9,11) equal to zero and taking into account that the sum in (9) consists of a single term, we obtain for the equilibrium density of a population monomorphic in *β*

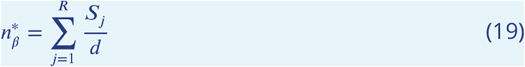

The invasion fitness of a rare mutant with uptake strategy *β*′ in a resident *β* at ecological equilibrium 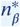 becomes:

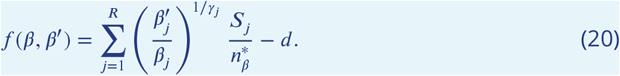

To derive the adaptive dynamics of *β*, we calculate the selection gradient *q*(*β*) = (*q*_1_(*β*), …, *g*_*R*_(*β*)) and project it on to the linear constraint space:

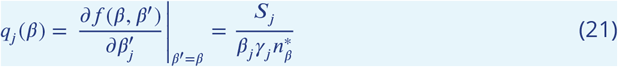

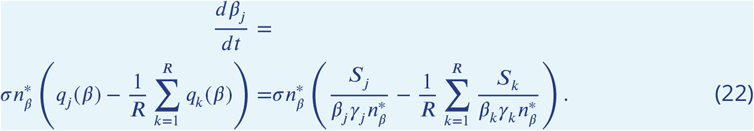

Here the term

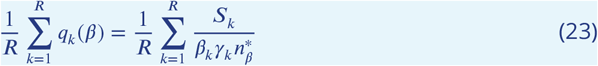

is the component of the selection gradient (21) that is orthogonal to the tradeoff hyperplane (note that 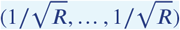 is a unit vector orthogonal to the tradeoff hyperplane).

If we set the mutational parameter *σ* = 1, the adaptive dynamics of *β* becomes:

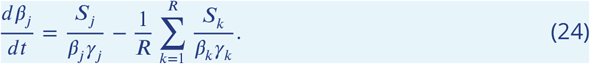

Note that we only need *R* − 1 equations due to the (linear) tradeoff. It is easy to see that (24) has a unique fixed point, i.e., there is a unique singular point for the adaptive dynamics given by:

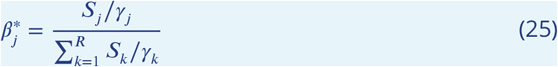

for *j* = 1, …, *R*. In terms of the original trait *α*, Eq. (25) is Eq. (8) in the main text.

To check for convergence stability of 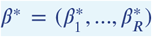, we have to calculate the Jacobian matrix *J* of the right hand side of (24), evaluated at the singular point *β*^∗^. It is easy to see that the *jk*-th element of *J* is

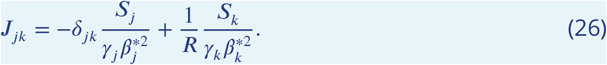

Thus, *J* is of the form

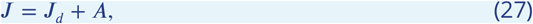

where *J*_*d*_ is a diagonal matrix with element 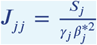, and *A* is a matrix whose elements in the *k*-th column are all identical and equal to 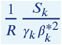. This implies that the matrix *A* maps any vector in phenotype space to a vector that is orthogonal to the tradeoff hyperplane (i.e., to a multiple of the vector (1, …, 1)). If Δ*β* = (Δ*β*_1_, …, Δ*β*_*R*_) is any vector of deviations from the singular point, it follows that the projection of *J*Δ*β* onto the tradeoff hyperplane is the same as the projection of the vector *J*_*d*_ Δ*β*. Since all eigenvalues of *J*_*d*_ are real and negative, it follows that the singular point *β*^∗^ is a local attractor, i.e., convergent stable, regardless of the exponents *γ*_*j*_, *j* = 1, …, *R*.

For evolutionary stability, we have to calculate the Hessian matrix *H* of second derivatives of the invasion fitness function, (20), with respect to the mutant trait *β*′ and evaluated at the singular trait value *β*^∗^. The *jk*-th element of *H* is

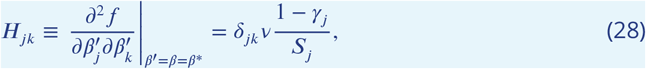

were *v* is a constant:

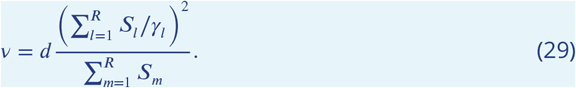

Thus, *H* is diagonal (due to the transformation from *α* to *β*), and *H* is negative definite, i.e., all eigenvalues are < 0, if and only if *γ*_*j*_ > 1 for all *j* = 1, …, *R*. Because the tradeoff hyperplane is linear in *β*, it follows that any index *j* with *γ*_*j*_ < 1 provides a branching direction *β*_*j*_, i.e., a direction in phenotype space along which evolutionary diversification is possible. More precisely, any direction in *β*-space (other than orthogonal to the tradeoff surface) along which the unconstrained Hessian (28) has a minimum corresponds to a direction on the tradeoff surface along which diversification is possible.

The results presented in the main text now follow from the above by setting *γ*_*j*_ = *γ* for *j* = 1, …, *R*. But the above analysis also suggests that with suitably chosen *γ*_*j*_, it is possible to generate evolutionary branching in some directions, but not in others. This is illustrated in Appendix 1 Figures 1 and 2. Our analysis and numerical procedure are applicable to an evolving system of populations with any number of resources. To facilitate visualization, in the following we consider just three resources, so that because of the constraint, each population is characterized by two independent parameters *α*_*i*_ Fig. 1 illustrates diversification in the direction of *α*_3_ (*γ*_3_ < 1) without diversification in the directions *α*_1_ and *α*_2_ (*γ*_1_, *γ*_2_ > 1).

**Appendix 1 Figure 2.**
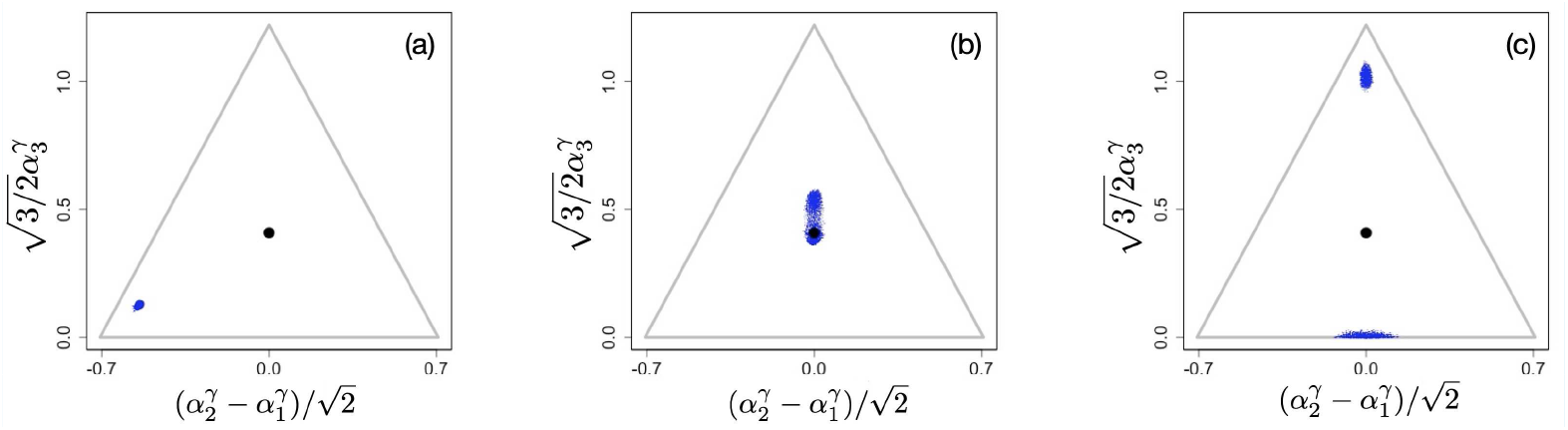
Example of evolutionary dynamics for *γ*_1_ = *γ*_2_ = 1.1 and *γ*_3_ = 0.9, showing convergence to the singular point and subsequent diversification only in the *α*_3_ direction. (Note that the dynamics are shown in the original *α*-phenotype space.) The corresponding video can be found **[here](https://figshare.com/s/3eb51bad58ae95b8b0f3)**, each frame in the video is separated by 2000 time steps. Other parameter values were *S*_*j*_ = 1, *μ*_*j*_ *K*_*j*_ = 0.1 for *j* = 1, 2, 3, and *d* = 0.25.

Fig. 1 illustrates diversification in the directions *α*_1_ and *α*_2_ (*γ*_1_, *γ*_2_ < 1), with no diversification in *α*_3_ (*γ*_3_ > 1).

**Appendix 1 Figure 3.**
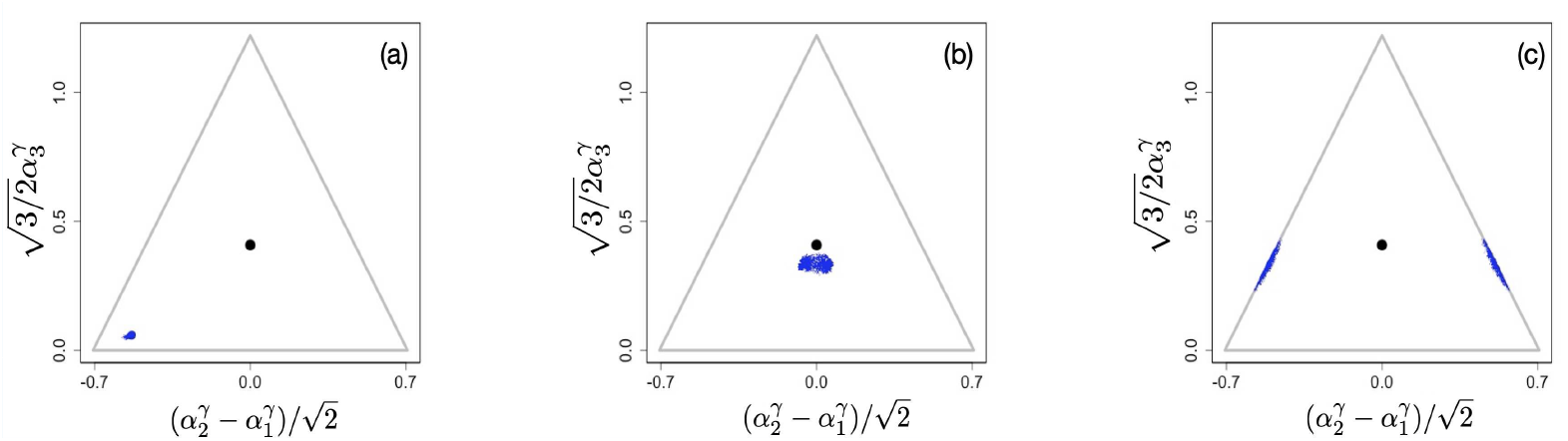
Example of evolutionary dynamics for *γ*_1_ = *γ*_2_ = 0.9 and *γ*_3_ = 1.2, showing convergence to the singular point and subsequent diversification only in *α*_1_ − *α*_2_ directions. (Note that the dynamics are shown in the original *α*-phenotype space.) The corresponding video can be found **[here](https://figshare.com/s/3eb51bad58ae95b8b0f3)**, each frame in the video is separated by 2000 time steps. Other parameter values were *S*_*j*_ = 1, *μ*_*j*_ *K*_*j*_ = 0.1 for *j* = 1, 2, 3, and *d* = 0.25.

#### Numerical Procedures

In the chemostat simulations, we numerically integrate the system of population dynamics equations (11) for *M* populations using a simple Euler update (*M* = 1 at the beginning of the simulations). After each integration step, the populations that fall below a small “extinction” threshold density (normally *n*_*min*_ = 10^−6^) are removed from the system. The resource concentrations *c*_*i*_ and uptake rates *r*_*i*_ are considered relaxed to their steady states for a given set of populations {*n*_*α*_},

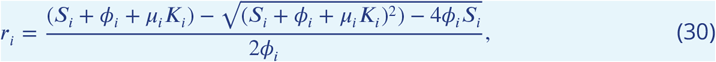

where *ϕ*_*i*_ = ∑_*α*_ *n*_*α*_ *α*_*i*_.

To simulate serial dilutions, we numerically integrate equations (6,7) for *M* populations and *R* resources using also the Euler update (*M* = 1 at the beginning of the simulations). After each integration step, the populations that fall below a small “extinction” threshold density (normally *n*_*min*_ = 10^−6^) are removed from the system. Once the resources are depleted so that the condition (8) is satisfied, the populations of all existing species are rescaled according to (5) and the resource concentrations are reset to *c*_*j*_ (0).

To mimic mutations in both simulation setups, periodically (typically once every Δ*t*_*mutation*_ = 1 time units) a mutant is split from an ancestor, which is randomly chosen with probability proportional to its total birth rate. The mutant’s phenotype is randomly offset from the ancestral phenotype along the constraint surface. The offset distance is drawn from a uniform distribution in the interval [−*m, m*]. Unless otherwise noted, *m* = 0.005. The mutant population is set to be 10% of the ancestral one, and the ancestor population is reduced by 10%. In addition to mutations, periodically (typically once every Δ*t*_*merge*_ = 100Δ*t*_*mutation*_ time units) populations that are within a distance *m* of each other are merged (preserving their phenotypic center of mass) and their population sizes added. Periodic repetition of mutation and merging procedures preserves the phenotypic variance necessary for evolution while limiting computational complexity.

This produces clouds or clusters of *α*-values in phenotype space (see figures), with each phenotype *α* representing a monomorphic population of individuals with that phenotype. Somewhat imprecisely, we refer to a distinct cluster of phenotypes as a species. The clusters move in phenotype space due to extinction and merging of phenotypes, and due to creation of new phenotypes by mutation. This movement represents evolution and occurs along the constraint surface. A diversification event occurs when a cluster corresponding to the diversifying species spontaneously splits into two or more clusters that diverge from each other and move apart.

#### Maintenance of diversity in ecological time scales

Here we briefly show how nonlinear trade-offs affect diversity on ecological time scale. To do this, we initiate the simulations with a set of e.g. randomly chosen phenotypes throughout phenotype space and then run the systems with the mutational process turned off. In Appendix 1 Fig. 3A we show the initial configuration used for three different scenarios with different exponents *γ* of the tradeoff (here we again assume that *γ*_*j*_ = *γ* for all *j*). The functional form of the tradeoffs has a profound effect on the number of surviving species, with many species coexisting when *γ* = 1, as reported in ***Posfai et al. (2017)***, (Appendix 1 Fig. 3B), but with typically only *R* species surviving when *γ* < 1 (Appendix 1 Fig. 3C) and only very few species surviving in the close vicinity of the singular point when *γ* > 1 (Appendix 1 Fig. 3D).

**Appendix 1 Figure 4.**
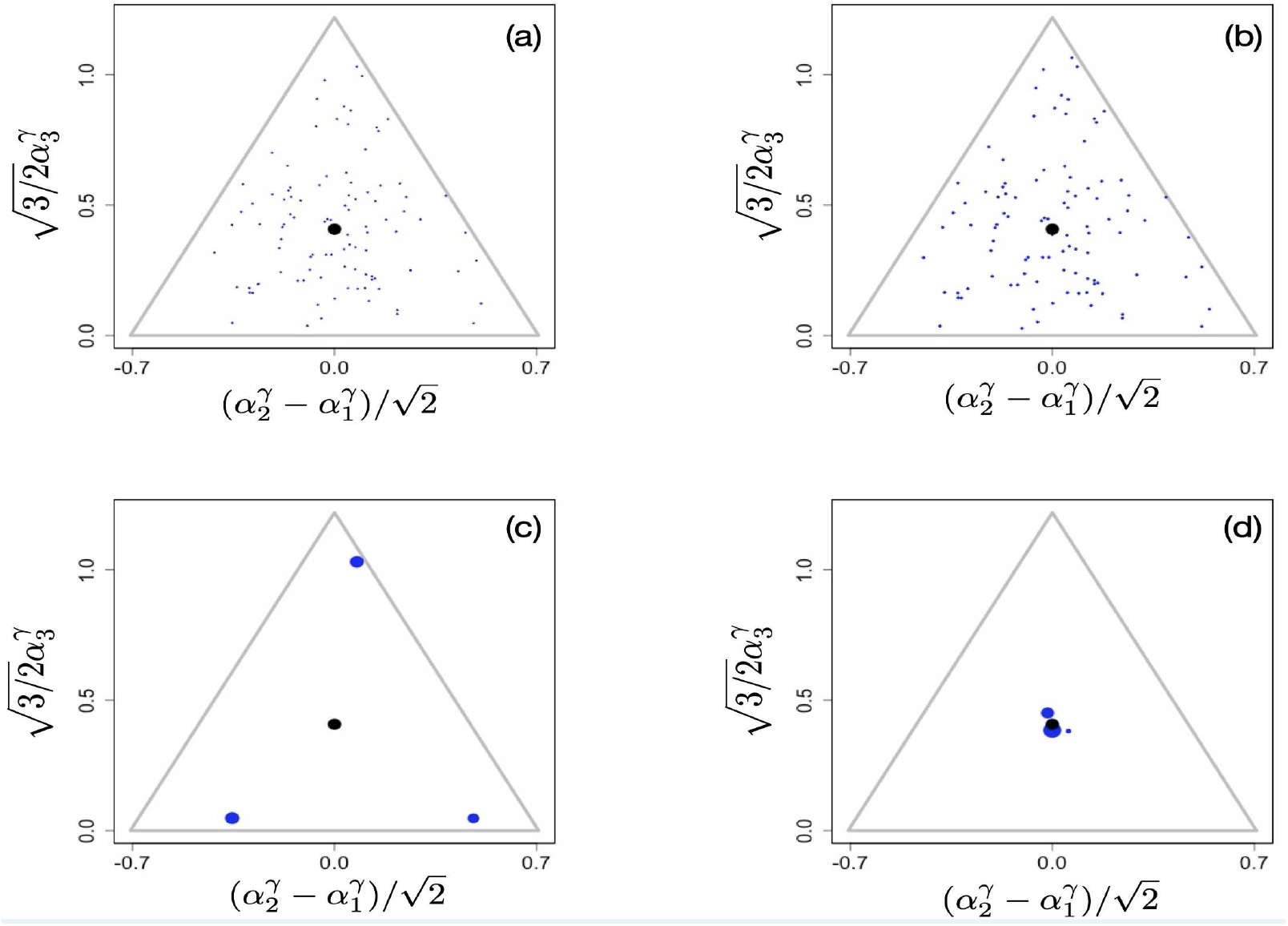
Initial population configuration of 100 randomly placed clusters in the phenotypic simplex (A), final configurations after 5000000 time units for *γ* = 1 (B), *γ* = 0.9 (C) and *γ* = 1.1 (D). Videos of the entire ecological processes can be found **[here](https://figshare.com/s/3eb51bad58ae95b8b0f3)**, time interval between frames increased as a geometric progression, *t*(*i* + 1) = 1.05*t*(*i*). Other parameter values were *S*_*j*_ = 1, *μ*_*j*_ *K*_*j*_ = 0.1 for *j* = 1, 2, 3, and *d* = 0.25.

Videos of ecological simulation of serial dilution scenario can be found **[here](https://figshare.com/s/3eb51bad58ae95b8b0f3)**

#### Nonlinear metabolic rates

Consider the consumption or transformation of a resource substance *c* into a downstream metabolic product *p* using a specific enzyme *α*:

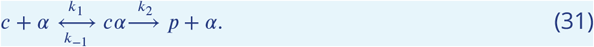

The assumption that the concentration of the complex *cα* instantaneously relaxes to its steady state defined by the current concentrations of *c* and *α* constitutes the MichaelisMenten approximation,

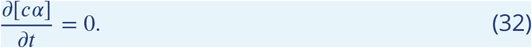

Assuming mass action kinetics and denoting by *c* and *α* the total (bound in the complex plus free) concentrations of the resource and enzyme, and dropping the traditional [] symbols for concentrations, one gets a quadratic equation for the the concentration *ψ* of the *cα* complex:

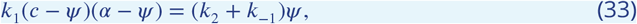

with the solution

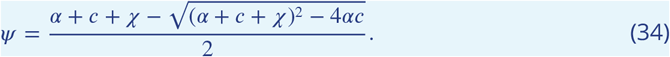

Here *χ* ≡ (*k*_−1_ +*k*_2_)/*k*_1_ is the dissociation constant for the complex. The more common form of the Michaelis-Menten approximation,

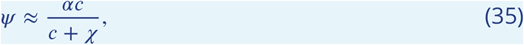

is obtained in the limit when the substrate concentration is much larger than that of the enzyme. (We note that this form is linear in *α* and is identical to the product of a Monod function and the corresponding *α* used in (11.)) Since

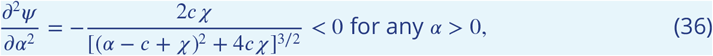

the resource uptake and the growth rates are always sub-linear in the concentration of the enzyme *α*.

## References

Abrams PA. Character displacement and niche shift analyzed using consumer-resource models of competition. Theoretical Population Biology. 1986; 29(1):107–160.

Ackermann M, Doebeli M. Evolution of niche width and adaptive diversification. Evolution. 2004; 58(12):2599–2612.

de Aguiar MAM, Baranger M, Baptestini EM, Kaufman L, Bar-Yam Y. Global patterns of speciation and diversity. Nature. 2009; 460(7253):384–387.

Archetti M, Scheuring I. Game theory of public goods in one-shot social dilemmas without assortment. Journal of theoretical biology. 2012; 299:9–20.

Bailey JE. Reflections on the scope and the future of metabolic engineering and its connections to functional genomics and drug discovery. Metabolic Engineering. 2001; 3(2):111–114.

Bolnick DI, Fitzpatrick BM. Sympatric Speciation: Models and Empirical Evidence. Annual Review of Ecology, Evolution, and Systematics. 2007; 38(1):459–487.

Borghans JA, De Boer RJ, Segel LA. Extending the quasi-steady state approximation by changing variables. Bulletin of mathematical biology. 1996; 58(1):43–63.

Bush GL. Sympatric speciation in animals: new wine in old bottles. Trends in Ecology & Evolution. 1994; 9(8):285–288.

Ciliberto A, Capuani F, Tyson JJ. Modeling networks of coupled enzymatic reactions using the total quasi-steady state approximation. PLoS Comput Biol. 2007; 3(3):e45.

Coyne JA, Orr HA. Speciation. Sinauer; 2004.

Coyne JA. Genetics and speciation. Nature. 1992; 355(6360):511–515.

Cui W, Marsland III R, Mehta P. Effect of resource dynamics on species packing in diverse ecosystems. Physical Review Letters. 2020; 125(4):048101.

Damore JA, Gore J. Understanding microbial cooperation. Journal of theoretical biology. 2012; 299:31–41.

Darwin C. On the Origin of Species by Means of Natural Selection. London: Murray; 1859. Or the Preservation of Favored Races in the Struggle for Life.

Dieckmann U, Doebeli M. On the origin of species by sympatric speciation. Nature. 1999; 400(6742):354–357.

Dieckmann U, Law R. The dynamical theory of coevolution: a derivation from stochastic ecological processes. Journal of Mathematical Biology. 1996; 34(5):579–612.

Doebeli M. Adaptive Diversification (MPB-48). Princeton University Press; 2011.

Doebeli M, Dieckmann U. Evolutionary Branching and Sympatric Speciation Caused by Different Types of Ecological Interactions. The American Naturalist. 2000; 156(S4):S77–S101.

Doebeli M, Dieckmann U. Speciation along environmental gradients. Nature. 2003; 421(6920):259–264.

Doebeli M, Ispolatov I. Continuously stable strategies as evolutionary branching points. Journal of Theoretical Biology. 2010; 266(4):529–535.

Doebeli M, Ispolatov I. Diversity and Coevolutionary Dynamics in High-Dimensional Phenotype Spaces. The American Naturalist. 2017; 189(2):105–120.

Erez A, Lopez JG, Weiner BG, Meir Y, Wingreen NS. Nutrient levels and trade-offs control diversity in a serial dilution ecosystem. eLife. 2020 sep; 9:e57790.

Friesen ML, Saxer G, Travisano M, Doebeli M. Experimental evidence for sympatric ecological diversification due to frequency-dependent competition in Scherichia coli. Evolution. 2004; 58(2):245–260.

Gavrilets S, Waxman D. Sympatric speciation by sexual conflict. Proceedings of the National Academy of Sciences. 2002 08; 99(16):10533.

Geritz SAH, Metz JA J, Kisdi E, Meszéna G. Dynamics of Adaptation and Evolutionary Branching. Phys Rev Lett. 1997 Mar; 78:2024–2027.

Goyal A, Maslov S. Diversity, Stability, and Reproducibility in Stochastically Assembled Microbial Ecosystems. Phys Rev Lett. 2018 Apr; 120:158102.

Guarner F, Malagelada JR. Gut flora in health and disease. The Lancet. 2003; 361(9356):512–519.

Hardin G. The Competitive Exclusion Principle. Science. 1960; 131(3409):1292–1297.

Hawksworth DL. The magnitude of fungal diversity: the 1.5 million species estimate revisited. Mycological Research. 2001; 105(12):1422–1432.

Hawksworth DL, Lücking R. 4. In: Fungal Diversity Revisited: 2.2 to 3.8 Million Species John Wiley & Sons, Ltd; 2017. p. 79–95.

Higashi M, Takimoto G, Yamamura N. Sympatric speciation by sexual selection. Nature. 1999; 402(6761):523–526.

Hill CM, Waightm RD, Bardsley WG. Does any enzyme follow the Michaelis—Menten equation? Molecular and cellular biochemistry. 1977; 15(3):173–178.

Hoefnagel MH, Starrenburg MJ, Martens DE, Hugenholtz J, Kleerebezem M, Van Swam II, Bongers R, Westerhoff HV, Snoep JL. Metabolic engineering of lactic acid bacteria, the combined approach: kinetic modelling, metabolic control and experimental analysisThe GenBank accession number for the sequence reported in this paper is AY046926. Microbiology. 2002; 148(4):1003–1013.

Hui C, Minoarivelo HO, Landi P. Modelling coevolution in ecological networks with adaptive dynamics. Mathematical Methods in the Applied Sciences. 2018; 41(18):8407–8422.

Hutchinson GE. The Paradox of the Plankton. The American Naturalist. 1961; 95(882):137–145.

Ispolatov I, Doebeli M. A note on the complexity of evolutionary dynamics in a classic consumer-resource model. Theoretical Ecology. 2020; 13(1):79–84.

Ito H, Sasaki A. Evolutionary branching under multi-dimensional evolutionary constraints. Journal of Theoretical Biology. 2016; 407:409–428.

Kisdi E. Trade-off geometries and the adaptive dynamics of two co-evolving species. Evolutionary Ecology Research. 2006; 8:959–973.

Kisdi É. Construction of multiple trade-offs to obtain arbitrary singularities of adaptive dynamics. Journal of Mathematical Biology. 2015; 70(5):1093–1117.

Larsen BB, Miller EC, Rhodes MK, Wiens JJ. Inordinate Fondness Multiplied and Redistributed: the Number of Species on Earth and the New Pie of Life. The Quarterly Review of Biology. 2017; 92(3):229–265.

Leimar O. Multidimensional convergence stability. Evolutionary Ecology Research. 2009; 11:191?208.

Lenski RE, Travisano M. Dynamics of adaptation and diversification: a 10,000-generation experiment with bacterial populations. Proceedings of the National Academy of Sciences. 1994; 91(15):6808–6814. https://www.pnas.org/content/91/15/6808, doi: 10.1073/pnas.91.15.6808.

Litchman E, Edwards KF, Klausmeier CA. Microbial resource utilization traits and trade-offs: implications for community structure, functioning, and biogeochemical impacts at present and in the future. Frontiers in Microbiology. 2015; 6:254.

Louca S, Doebeli M. Calibration and analysis of genome-based models for microbial ecology. eLife. 2015 oct; 4:e08208.

Louca S, Doebeli M. Calibration and analysis of genome-based models for microbial ecology. eLife. 2015; 4.

Ma J, Levin SA. The Evolution of Resource Adaptation: How Generalist and Specialist Consumers Evolve. Bulletin of Mathematical Biology. 2006; 68(5):1111–1123.

Macarthur R, Levins R. The Limiting Similarity, Convergence, and Divergence of Coexisting Species. The American Naturalist. 1967; 101(921):377–385.

Marianayagam NJ, Sunde M, Matthews JM. The power of two: protein dimerization in biology. Trends in biochemical sciences. 2004; 29(11):618–625.

May RM. How Many Species Are There on Earth? Science. 1988; 241(4872):1441–1449.

Mayr E, Mayr AAPZMCZE. Animal Species and Evolution. Belknap Press, Belknap Press of Harvard University Press; 1963.

Metz JA J, Nisbet RM, Geritz SAH. How should we define ‘fitness’ for general ecological scenarios? Trends in Ecology & Evolution. 1992; 7(6):198–202.

Nelson David L (David Lee). Lehninger principles of biochemistry. Fourth edition. New York : W.H. Freeman, 2005.; 2005.

Pacciani-Mori L, Giometto A, Suweis S, Maritan A. Dynamic metabolic adaptation can promote species coexistence in competitive communities. PLoS computational biology. 2020; 16(5):e1007896.

Pacciani-Mori L, Giometto A, Suweis S, Maritan A. Dynamic metabolic adaptation can promote species coexistence in competitive microbial communities. PLOS Computational Biology. 2020 05; 16(5):1–18.

Posfai A, Taillefumier T, Wingreen NS. Metabolic Trade-Offs Promote Diversity in a Model Ecosystem. Phys Rev Lett. 2017 Jan; 118:028103.

Ricard J, Noat G. Catalytic efficiency, kinetic co-operativity of oligomeric enzymes and evolution. Journal of theoretical biology. 1986; 123(4):431–451.

Rice WR, Hostert EE. Laboratory Experiments of speciation: What have we learned in 40 years? Evolution. 1993; 47(6):1637–1653.

Schmitt DL, An S. Spatial organization of metabolic enzyme complexes in cells. Biochemistry. 2017; 56(25):3184–3196.

Shoresh N, Hegreness M, Kishony R. Evolution exacerbates the paradox of the plankton. Proceedings of the National Academy of Sciences. 2008; 105(34):12365–12369.

Steel M, Penny D. Common ancestry put to the test. Nature. 2010; 465(7295):168–169.

Theobald DL. A formal test of the theory of universal common ancestry. Nature. 2010; 465(7295):219–222.

Weiner BG, Posfai A, Wingreen NS. Spatial ecology of territorial populations. Proceedings of the National Academy of Sciences. 2019; 116(36):17874–17879.

